# Genetic variation for ontogenetic shifts in metabolism underlies physiological homeostasis in *Drosophila*

**DOI:** 10.1101/456269

**Authors:** Omera B. Matoo, Cole R. Julick, Kristi L. Montooth

## Abstract

Organismal physiology emerges from metabolic pathways and structures that can vary across development and among individuals. Here we tested whether genetic variation at one level of physiology can be buffered at higher levels during development by the inherent capacity for homeostasis in physiological systems. We found that the fundamental scaling relationship between mass and metabolic rate, as well as the oxidative capacity per mitochondria, differed significantly across development in the fruit fly *Drosophila*. However, mitochondrial respiration rate was maintained across development at similar levels. Furthermore, genotypes clustered into two types—those that switched to aerobic, mitochondrial ATP production before the second instar and those that relied on anaerobic production of ATP via glycolysis through the second instar. Despite genetic variation for the timing of this metabolic shift, second-instar metabolic rate was more robust to genetic variation than was the metabolic rate of other instars. We also found that a mitochondrial-nuclear genotype with disrupted mitochondrial function both increased aerobic capacity more through development and relied more heavily on anaerobic ATP production relative to wildtype genotypes. By taking advantage of both ways of making ATP, this genotype maintained mitochondrial respiratory capacity, but also generated more free radicals and had decreased mitochondrial membrane potential, potentially as a physiological-defense mechanism. Taken together, the data revealed that genetic defects in core physiology can be buffered at the organismal level via physiological compensation and that natural populations likely harbor genetic variation for distinct metabolic strategies in development that generate similar organismal outcomes.

## Introduction

Metabolism is the sum total of biochemical processes by which organisms use energy to sustain life and fuel reproduction, and an individual's metabolic rate is often interpreted as an integrated measure of its “pace of life”(Glazier, 2005, 2014, 2015). Early surveys of molecular variation revealed a surprising amount of genetic variation segregating in natural populations at the loci encoding these biochemical processes (Harris, 1966; Hubby and Lewontin, 1966; Lewontin and Hubby, 1966), a pattern that has been historically used to advocate both for the predominance of classical mutation, drift and purifying selection forces (Kimura 1983) and for the maintenance of variation through selection (Gillespie, 1999) (reviewed by Charlesworth and Charlesworth, 2016). Subsequent surveys revealed substantial quantitative genetic variation in metabolic enzyme activities within species, arising from both molecular variation at enzyme-encoding loci, as well as trans-acting and epistatic variation throughout the genome (Clark and Wang, 1997; Clark *et al.*, 1995a, 1995b; Laurie-Ahlberg *et al.*, 1980, 1982; Mitchell-Olds and Pedersen, 1998; Montooth *et al.*, 2003). In a few cases, this biochemical variation has been linked to variation at higher levels of physiological or organismal performance (Crawford and Oleksiak, 2007; Laurie-Ahlberg *et al.*, 1982; Montooth *et al.*, 2003; Watt *et al.*, 1983) and in some cases may be adaptive (Tishkoff *et al.*, 2001; Verrelli and Eanes, 2001; Watt, 1977; Watt *et al.*, 2003). However, we still lack a general mechanistic understanding of how genetic variation in the pathways of metabolism is transformed up the hierarchical levels of biological organization to result in variation for the organismal performance traits that determine fitness. This understanding is important for consideration of metabolism as an adaptive phenotype and for predicting how selection on metabolic performance will shape variation in genomes.

A challenge to connecting genetic variation in biochemical processes to metabolic performance, is that metabolism is an emergent property of interacting biochemical, structural, regulatory and physiological systems, often arranged in hierarchical functional modules (Barabási and Oltvai, 2004; Jeong *et al.*, 2000; Ravasz *et al.*, 2002; Strogatz, 2001). In addition, metabolic enzymes and metabolites have potential “moonlighting” functions in the signaling that underlies metabolic homeostasis (Marden, 2013; Boukouris *et al.*, 2016). The capacity for homeostasis in physiological systems, also suggests that genetic variation in biochemical processes at one level of the energetic hierarchy may not necessarily result in organismal fitness variation. In other words, the physiological regulatory processes that maintain energy homeostasis may provide stability in metabolic trajectories, in an analogous way to the canalized developmental trajectories envisioned by Waddington (Meiklejohn and Hartl, 2002; Waddington, 1942, 1957). Furthermore, a diversity of biochemical pathways may be available to achieve similar energetic outputs. Finally, the hierarchical biological processes that contribute to metabolism are influenced by both extrinsic (e.g., temperature, resource availability, habitat, and infection status) and intrinsic (e.g., genotype, life stage, sex, activity level, and reproductive status) factors [reviewed (Glazier, 2005)], such that genetic variation in biochemical processes may affect organismal performance and fitness in only a subset of conditions. Such conditionally neutral variation is expected to experience less effective selection, as it will be seen by selection in only a fraction of contexts (Van Dyken and Wade, 2010).

Development is a potentially important context for the expression of genetic variation in metabolism. During development, organisms partition energetic resources between the competing demands of growth, development, maintenance, and storage for future reproduction. Energy homeostasis during development is largely achieved by feedback controls where energy-demand processes increase the concentration of ADP, which can then be fed as a substrate for energy-supply processes to generate ATP. As key metabolic organelles, the mitochondria play a central role in the energy supply-demand balance. Not only the abundance and activity of mitochondria, but also the surface area, membrane composition, and cellular network structure of mitochondria have been reported to affect metabolism (Miettinen and Björklund, 2017; Porter and Brand, 1993; Porter *et al.*, 1996). In addition, both mitochondrial and nuclear genomes interact closely to form the protein complexes of oxidative phosphorylation (OXPHOS) that drive aerobic ATP production, creating the potential for inter-genomic epistasis to underlie variation in metabolic phenotypes. At present, our understanding of the how the underlying genetic architecture and its variation affects metabolism is incomplete; but studies indicate that both nuclear DNA (nDNA) (Montooth et al., 2003; Nespolo *et al.*, 2007; Tieleman *et al.*, 2009), mitochondrial DNA (mtDNA) (Arnqvist *et al.*, 2010; Ballard and Rand, 2005; Kurbalija *et al.*, 2014; Martin, 1995), and interactions between genomes and environment are involved in determining metabolism (Hoekstra *et al.*, 2013; Hoekstra *et al.*, 2018).

Energy balance is particularly keen in holometabolous species where larval development and growth tends to be rapid and massive, requiring simultaneous accumulation of the resources needed to fuel metamorphosis and emergence as a reproductive adult. *Drosophila melanogaster* is an especially powerful system to study developmental metabolism, given the genetic resources and an approximately 200-fold increase in body mass across three larval instars (Church and Robertson, 1966). There is evidence of significant genetic variation for metabolic rate within *Drosophila* species (Montooth *et al.*, 2003; Hoekstra *et al.*, 2013), and mitochondrial–nuclear genotypes that disrupt mitochondrial function also adversely affect larval metabolic and development rates (Meiklejohn *et al.*, 2013; Hoekstra *et al.*, 2013). Transcriptomic and metabolic profiling in *D. melanogaster* indicate the dynamic nature of energy homeostasis that draws on both aerobic and anaerobic energy production, as well as the presence of proliferative metabolic programs during larval development (Graveley *et al.*, 2011; Tennessen *et al.*, 2011). Despite this wealth of data, at present, we lack a detailed understanding of the links between genome variation, mitochondrial function and organismal metabolic rate during development in *Drosophila*.

In the present study, we tested whether metabolic strategies in *D. melanogaster* varied among genotypes and across larval instars for both wildtype and mitochondrial-nuclear genotypes that generate energetic inefficiencies. We found that there is significant variation for the ontogeny of metabolism at the level of mitochondrial aerobic capacity, but that this variation can be buffered at higher levels of metabolic performance via physiological homeostasis. In this way, we show that there may be multiple genotypic and physiological paths to equivalent organismal outcomes within populations.

## Methods

### Drosophila stocks and maintenance

We used four *Drosophila* mitochondrial-nuclear genotypes generated by Montooth *et al.*, 2010. The (*mtDNA*);*nuclear* genotype (*simw*^501^);*OreR* has a genetic incompatibility that decreases oxidative phosphorylation (OXPHOS) activity putatively via compromised mitochondrial protein translation, and disrupts larval metabolic rate, resulting in delayed development, decreased immune function, and reduced female fecundity (Meiklejohn *et al.*, 2013; Hoekstra *et al.*, 2013; Holmbeck *et al.*, 2015; Zhang *et al.*, 2017; Hoekstra *et al.*, 2018; Buchanan *et al.*, 2018). This mitochondrial-nuclear (hereafter referred to as mito-nuclear) incompatibility arises from an epistatic interaction between naturally-occurring single nucleotide polymorphisms (SNPs) in the mt-tRNA^Tyr^ gene and the nuclear-encoded mt-tyrosyl-tRNA synthetase gene *Aatm* that aminoacylates this mitochondrial tRNA (Meiklejohn *et al.*, 2013). The other three genotypes – (*ore*);*OreR*, (*simw*^501^);*Aut*, and (*ore*);*Aut* – serve as wildtype genetic controls that enabled us to test for the effects of mitochondrial and nuclear genotypes, separately and interactively, on developmental physiology. Additionally, we measured traits in two inbred genotypes sampled in Vermont (*VT4* and *VT10*) as representatives of natural populations that were not manipulated to generate specific mito-nuclear genotypes.

All genotypes were raised on standard cornmeal-molasses-yeast *Drosophila* media and acclimated to 25°C with a 12 h:12 h dark:light cycle for at least three generations prior to all experiments. To collect first-, second- and third-instar larvae, adults were allowed to lay eggs for 3-4 hours on standard media, and larvae from these cohorts were staged based on developmental time and distinguishing morphological features.

### Larval metabolic rate

Routine metabolic rate was measured as the rate of CO_2_ produced by groups of twenty larvae of the same instar and genotype using a flow-through respirometry system (Sable Systems International, Henderson, NV) with established protocols (Hoekstra *et al.*, 2013). Groups of larvae were collected onto the cap of 1.7 mL tube containing 0.5 mL of fly media and placed inside one of four respirometry chambers that were housed in a temperature-controlled cabinet (Tritech™ Research, Los Angeles, CA) maintained at 25°C. Between 8 and 13 biological replicates for each genotype and instar were randomized across chambers and respirometry runs, during which each group of larvae was sampled for CO_2_ production for two, ten-minute periods. CO_2_ that accumulated in the chambers as a result of larval metabolism was detected using an infrared CO_2_ analyzer (Li-Cor 7000 CO_2_/H_2_O Analyzer; LI-COR, Lincoln, NE). 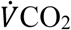 was calculated from the mean fractional increase in CO_2_ at a constant air-flow rate of 100 ml/min over a 10 min time interval for each replicate after baseline drift correction. The wet weight of the group of larvae was recorded using a Cubis^®^ microbalance (Sartorius AG, Göttingen, Germany) at the beginning of each respirometry run.

### Isolation of Mitochondria

Mitochondria were isolated from larvae following a protocol modified from Aw *et al.*, 2016. 100-200 larvae were collected and rinsed with larval wash buffer (0.7% NaCl and 0.1 % Triton X-100). Larvae were very gently homogenized in 300-500 µL of chilled isolation buffer (154 mM KCl, 1 mM EDTA, pH 7.4) in a glass-teflon Thomas^®^ homogenizer on ice. The homogenate was filtered through a nylon cloth into a clean chilled microcentrifuge tube. The homogenate was then centrifuged at 1500g for 8 min at 4°C (Eppendorf Centrifuge 5810R). The resulting mitochondrial pellet was suspended in 40-50 µL of ice-cold mitochondrial assay solution (MAS: 15 mM KCl, 10 mM KH_2_PO_4_, 2 mM MgCl_2_, 3 mM HEPES, 1 mM EGTA, FA-free BSA 0.2%, pH 7.2). Unless otherwise stated, all chemicals were purchased from Sigma Aldrich (St Louis, MO) or Fisher Scientific (Pittsburgh, PA) and were of reagent grade or higher.

### Mitochondrial Respiration

Oxygen consumption of freshly isolated mitochondria was measured using the Oxygraph Plus System (Hansatech Instruments, Norfolk, UK) in 3 mL water jacketed glass chambers equipped with a magnetic stirrer and Clark-type oxygen electrodes. Temperature of the respiration chambers was kept constant at 25°C using a Fisher Isotemp^®^ 4100 R20 refrigerated water circulator (Fisher Scientific, Hampton, NH). A two-point calibration of electrodes using air-saturated distilled water and sodium sulfite was done for establishing 100% and zero oxygen levels in the chamber, respectively. The assay was completed within 2 hours of mitochondrial isolation, and six or seven biological replicates were measured for each larval stage of each genotype. 50 µL (~1.5 mg protein) of mitochondrial suspension was added to 950 µL of MAS in the respiration chamber. Pyruvate (5mM) and malate (2.5mM) were used as respiratory substrates at saturating amounts. Maximum respiration (State 3) was achieved by adding 400 µM of ADP, and State 4^+^ respiration was calculated as described by Chance and Williams 1955 by adding 2.5 µg ml^-1^ oligomycin. Oligomycin is an ATPase inhibitor and State 4^+^ gives an estimate of oxygen consumption linked to mitochondrial proton leak, rather than to ATP production, at high membrane potential (Brand *et al.*, 1994). Uncoupled respiration (State 3u), indicative of maximum respiration or electron transport system (ETS) capacity, is achieved by adding 0.5 µM of carbonyl cyanide m-chlorophenyl hydrazone (CCCP). CCCP is a protonophore that increases proton permeability in mitochondria and effectively disconnects ETS from ATPase. Data were acquired and respiration rates were corrected for electrode drift using the OxyTrace^+^ software. The respiratory control ratio (RCR) was calculated as the ratio of State 3 over State 4^+^ (Estabrook 1967). Respiration rates were normalized by unit mitochondrial protein added. Protein concentrations were determined using Bio-Rad Protein Assay Dye Reagent Concentrate (Bio-Rad, 5000006) and bovine serum albumin (BSA) as a standard.

### Mitochondrial membrane potential (ΔΨ_m_)

Mitochondrial membrane potential was measured using the JC-1 indicator dye (Fisher Scientific, Hampton, NH) following a protocol modified from Villa-Cuesta *et al.*, 2014.100 mg of larvae were weighed and used to isolate mitochondria as described above. Approximately 1.4 mg of mitochondrial protein was added and the final volume was increased to 300 µL using MAS. 3 µL of a 1 µg/µL solution of JC-1 dissolved in dimethyl sulfoxide (DMSO) was added to the suspension. Mitochondrial samples were incubated for 30 min at 37°C protected from light. At the end of incubation, samples were centrifuged for 3 min at 6000g and suspended in 600 µL of fresh MAS. Mitochondrial membrane potential was expressed as the ratio of fluorescence for aggregate:monomeric forms of JC-1 at red (excitation 485 nm, emission 600 nm) and green (excitation 485 nm, emission 530 nm) wavelengths respectively. 50 µM of CCCP was added to collapse membrane potential as a negative control.

### Citrate synthase activity

Citrate synthase activity was measured following the protocol from Meiklejohn *et al.*, 2013. 100-200 larvae were homogenized in 1 mL chilled isolation buffer (225 mM mannitol, 75 mM sucrose, 10 mM MOPS, 1 mM EGTA, 0.5% fatty acid-free BSA, pH 7.2) using a glass-teflon Thomas^®^ homogenizer. The homogenate was centrifuged at 300g for 5 min at 4°C (Eppendorf Centrifuge 5810R). The supernatant was transferred into a clean tube and centrifuged again at 6000g for 10 min at 4°C. The resulting mitochondrial pellet was resuspended in 50 µL of respiration buffer (225 mM mannitol, 75 mM sucrose, 10 mM KCl, 10 mM Tris-HCl, and 5 mM KH_2_PO_4_, pH 7.2). All samples were stored at -80°C till further analysis.

Maximum citrate synthase activity (V_max_) of the mitochondrial extracts was measured spectrophotometrically at 30°C using a Synergy 2 plate reader (BioTek, VT, USA). 6 µg of mitochondrial protein was added to the assay mixture containing 100mM Tris-HCl (pH 8.0), 2.5 mM EDTA, 100 µM Acetyl Co-A and 100 µM of DTNB [5,5′-dithiobis (2-nitrobenzoic acid)]. The reaction was monitored for 2 min as a background reading. The reaction was then started by adding 500 µM oxaloacetate to the assay to generate CoA-SH. CoA-SH is then detected by its reaction with DTNB to form a yellow product (mercaptide ion) that was measured using absorbance at 412 nm. Enzyme activity was normalized by protein concentration of the sample added. Six biological samples per genotype and instar were measured, each with two technical replicates.

### Lactate quantification

Whole-body lactate concentrations were measured by an NAD^+^/NADH-linked fluorescent assay following the protocol of Callier *et al.*, 2015. 100-200 larvae were homogenized in 100-500 µL of 17.5% perchloric acid and centrifuged at 14,000g for 2 min at 4°C (Eppendorf Centrifuge 5810R). Following precipitation of proteins, the clear supernatant was transferred into a clean tube and neutralized with a buffer containing 2 M of KOH and 0.3 M of MOPS, and again centrifuged at 14,000g for 2 min at 4°C. 20-50 µL of neutralized sample was added to the assay buffer (pH 9.5) containing a final concentration of 1000 mM hydrazine, 100 mM Tris-base, 1.4 mM EDTA and 2.5 mM NAD^+^ in a 96-well plate (Micro Flour^®^ 1). The assay was performed in fluorescence mode (Ex/Em = 360/460 nm) using a Synergy H1 Hybrid Reader (BioTek, VT, USA). After incubating the plate at 5 min at room temperature, a background reading was taken. 17.5 U/well lactate dehydrogenase (Sigma L3916) diluted with Tris buffer was then added to each sample and the reaction mixture was allowed to incubate at 37°C for 30 min protected from light. A second reading was then taken to measure NADH levels, after correcting for background fluorescence. Six biological samples per genotype and instar were measured, each with two technical replicates. Sodium lactate was used as a standard for the assay. Lactate concentrations in the samples were normalized by wet weight of the larvae.

### Hydrogen peroxide (H_2_O_2_) quantification

100-200 larvae were weighed, rinsed with larval wash buffer (0.7% NaCl and 0.1 % Triton X100) and homogenized in 500 µL of pre-chilled assay buffer (pH 7.5) containing 20 mM HEPES, 100 mM KCl, 5% glycerol, 10 mM EDTA, 0.1% Triton X-100, 1 mM PMSF (Sigma P7626) and 1:10 (v/v) protease inhibitor cocktail (Sigma P2714) using a glass-teflon Thomas^®^ homogenizer. The homogenate was centrifuged at 200g for 5 min at 4°C, and the supernatant was stored at -80°C. H_2_O_2_ concentration was determined with a fluorometric Hydrogen Peroxide (H_2_O_2_) Assay Kit (Sigma MAK 165) following the manufacturer’s protocol in a 96-well plate (Micro Flour^®^ 1) using the Synergy H1 Hybrid Reader (BioTek, VT, USA). Six biological samples per genotype and instar were measured, each with two technical replicates. H_2_O_2_ concentrations in the samples were expressed as nM/µg of protein.

### Statistical analyses

All statistical analyses used the statistical package R version 2.15.1 (R Development Core Team 2011). We implemented standard major-axis regression in the R-package SMATR (Warton *et al.*, 2006; Hoekstra *et al.*, 2013) to estimate the relationship between log-transformed mass and 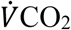, and to test for larval-instar and genetic effects on the slope of this relationship. When there was statistical evidence for a common slope among genotypes, we fit the common slope to test for effects of genotype on the y-intercept (i.e., genetic effects on the mass-specific metabolic rate). We removed a single observation where a first-instar replicate had a 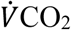 value less than zero. Analysis of variance (ANOVA) was used to test for the fixed effects of mtDNA, nuclear genome, larval instar, and all interactions on lactate accumulation, H_2_O_2_ concentration, mitochondrial physiology (State3, State4, uncoupled respiration, RCR^+^, and ΔΨ_m_) and citrate synthase activity in an ANOVA design. Post-hoc comparisons among instars within genotypes and among genotypes within instar were evaluated using Tukey HSD tests.

### Data Availability

Stocks and strains are available upon request. Supplemental files are available at FigShare. Phenotypic data will be deposited in the Dryad Digital Repository upon publication.

## Results

### Metabolic rate scaling with mass varies across larval instars and genotypes

Metabolic rate scales with mass according to the power function *R* = *aM^b^*, where *a* is the constant scaling coefficient, *M* is mass, and *b* is the scaling exponent. The scaling exponent *b*, estimated by the slope of the relationship between log-transformed metabolic rate and mass, differed significantly across larval instars (Fig. 1A) (*LR* = 18.1, *df* = 2, *P* = 0.0001). Metabolic scaling with body mass was hypermetric in first-instar larvae (*b* (CI) = 1.42 (1.21, 1.67)), isometric in second-instar larvae (*b* = 1.04 (0.95, 1.15)), and hypometric in third-instar larvae (*b* = 0.85 (0.71, 1.01)). Within first- and second-instar larvae, there was no evidence that metabolic scaling with mass differed significantly among genotypes, nor were there significant effects of genotype on the elevation of the fitted relationship (i.e., on the mass-specific metabolic rate) (Fig. 1B, 1C and Table 1). However, there was more variance among genotypes in mass-specific metabolic rate in first-instar larvae relative to second-instar larvae (Fig. 1B, 1C and Table 1). Metabolic scaling with mass in third-instar larvae differed significantly among genotypes, as evidenced by significantly different slopes (Fig. 1D and Table 1). The variation in metabolic scaling with mass did not result from natural lines differing from mito-nuclear genotypes, but rather from variation in the scaling exponent within both sets of genotypes. The pattern was significant regardless of the inclusion of several data points that, while not statistical outliers, did appear as outliers in the relationship between metabolic rate and mass (Fig. 1D and Supplemental Table 1).

**Figure 1.**
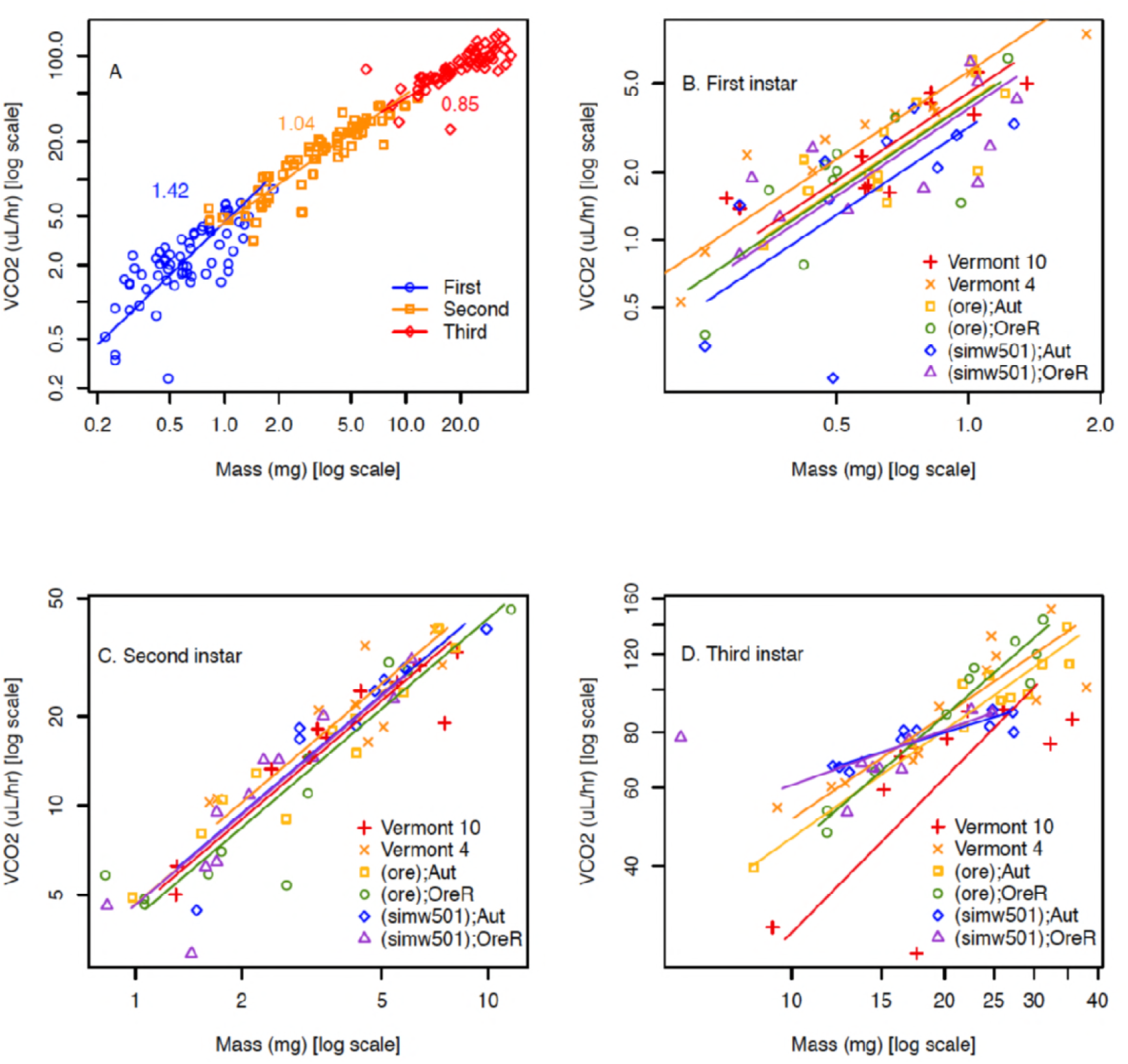
Larval metabolic rate. Metabolic scaling with mass varied across larval development and among genotypes. A) The mass-scaling exponent for routine metabolic rate 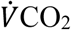 differed significantly among instars (*LR* = 18.1, *df* = 2, *P* = 0.0001), with the relationship between metabolic rate and mass becoming more shallow across development. B,C) There was more genetic variation for metabolic rate in first-instar larvae, relative to second-instar larvae. D) Mass-scaling exponents differed significantly among genotypes in the third instar of development (Table 1 and Supplemental Table 1).

**Table 1.** Genetic effects on routine metabolic rate (RMR) as a function of mass across developmental instars

| Phenotype | Genotype | Slope (95% CI)<sup>1</sup> | Y-intercept<sup>2</sup> |
| --- | --- | --- | --- |
| First-instar RMR |  | H<sub>0</sub>: equal slopes (LR=4.61, df=5, P=0.46) | H<sub>0</sub>: no elevation difference (Wald=9.28, df=5, P=0.10) |
|  | Common slope | 1.29 (1.10, 1.54) |  |
|  | VT10 |  | 0.66 (0.54, 0.77) |
|  | VT4 |  | 0.75 (0.64, 0.86) |
|  | (ore);Aut |  | 0.62 (0.51, 0.73) |
|  | (ore);OreR |  | 0.61 (0.43, 0.79) |
|  | (simw<sup>501</sup>);Aut |  | 0.50 (0.26, 0.75) |
|  | (simw<sup>501</sup>);OreR |  | 0.59 (0.42, 0.75) |
| Second-instar RMR |  | H<sub>0</sub>: equal slopes (LR=3.99, df=5, P=0.55) | H<sub>0</sub>: no elevation difference (Wald=2.58, df=5, P=0.76) |
|  | Common slope | 1.01 (0.90, 1.13) |  |
|  | VT10 |  | 0.65 (0.55, 0.75) |
|  | VT4 |  | 0.71 (0.6, 0.81) |
|  | (ore);Aut |  | 0.67 (0.58, 0.75) |
|  | (ore);OreR |  | 0.62 (0.51, 0.73) |
|  | (simw<sup>501</sup>);Aut |  | 0.67 (0.55, 0.79) |
|  | (simw<sup>501</sup>);OreR |  | 0.67 (0.57, 0.76) |
| Third-instar RMR |  | H<sub>0</sub>: equal slopes (LR=20.1, df=5, P=0.001) |  |
|  | VT10 | 1.16 (0.65, 2.06) |  |
|  | VT4 | 0.78 (0.54, 1.10) |  |
|  | (ore);Aut | 0.81 (0.65, 1.00) |  |
|  | (ore);OreR | 1.00 (0.81, 1.24) |  |
|  | (simw<sup>501</sup>);Aut | 0.36 (0.25, 0.52) |  |
|  | (simw<sup>501</sup>);OreR | 0.41 (0.19, 0.89) |  |

<sup>1</sup> Either a common or genotype-specific slope with confidence interval, when justified by the test for equal slopes among genotypes.
<sup>2</sup> In no case was there evidence that there was a shift in mass along the x-axis among genotypes ( $P > 0.09$ ).

### Mitochondrial respiration is similar across larval instars and genotypes

Despite these ontogenetic and genetic differences in the scaling of organismal metabolic rate with mass, second- and third-instar larvae had similar rates of mitochondrial oxygen consumption linked to ATP production (i.e., State 3 respiration) per unit of mitochondrial protein in both mito-nuclear genotypes (*instar*, *P =* 0.13) and natural genotypes (*instar*, *P =* 0.12) (Fig. 2) (Supplemental Table 2). State 3 respiration from first-instar larvae mitochondrial preparations were either below our detection limits or of low-quality, even when including similar amounts of larval mass in the preparation. This indicates that there is likely an increase in mitochondrial quantity or functional capacity between the first- and second-larval instars. Furthermore, State 3 respiration did not differ significantly among mito-nuclear or natural genotypes, nor were there any significant interactions between instar and genetic factors (Supplemental Table 2). Maximum respiratory capacity of mitochondria (or CCCP- induced uncoupled respiration) was also maintained across instars in all mito-nuclear genotypes (*instar*, *P*= 0.18) (Supplemental Fig. 1A and Supplemental Table 2). However, the natural genotype *VT10* had a significantly elevated maximal respiratory capacity in the second instar that resulted in a significant instar-by-genotype interaction (*P*= 0.001) (Supplemental Fig. 1A and Supplemental Table 2).

**Figure 2.**
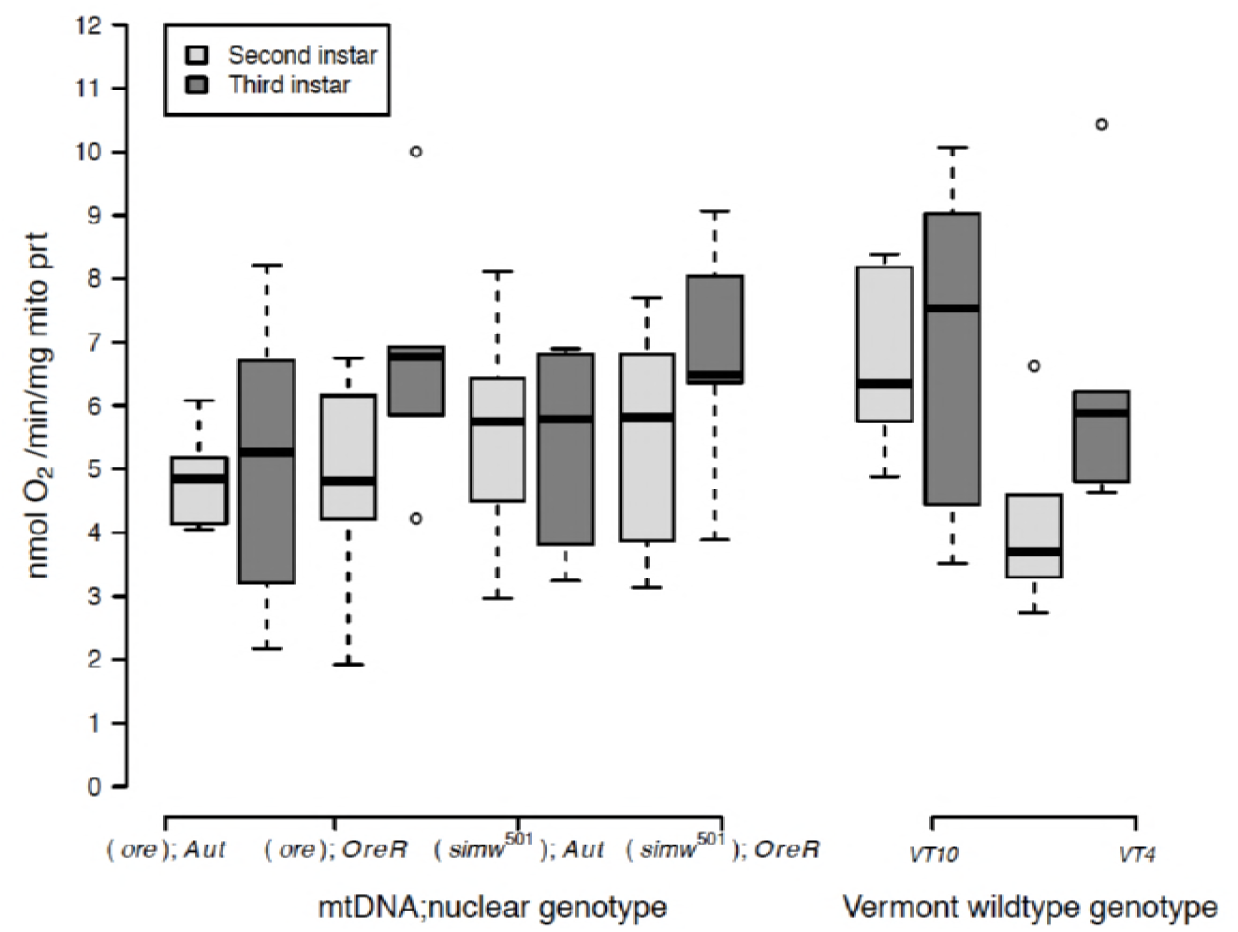
State 3 mitochondrial O_2_ consumption. Oxygen-coupled ATP production, measured by the State 3 mitochondrial oxygen consumption per unit of mitochondrial protein, was maintained at statistically similar levels across genotypes and instars (Supplemental Table 2).

Healthy mitochondria have high rates of oxygen consumption and ATP production when ADP is abundant (i.e., State 3 respiration), but low rates of oxygen consumption in the absence of ATP synthesis (i.e., State 4^+^ respiration). The ratio of these two measures is called the respiratory control ratio (RCR^+^). While the RCR^+^ was generally maintained at a ratio of 2-3 across genotypes and instars, two genotypes, (*ore*);*OreR* and *VT10,* had elevated RCR in third-instar larvae that contributed to a significant instar-by-genotype interaction in both mito-nuclear (instar x nuclear, *P* = 0.004) and natural genotypes (instar x genotype, *P* = 0.0001) (Supplemental Fig. 1B and Supplemental Table 2). This was due to decreased State 4^+^ respiration in second-instar mitochondria from these genotypes (Supplemental Fig. 1C and Supplemental Table 2).

### Certain genotypes use anaerobic ATP production further into development

We measured the activity of citrate synthase, a nuclear-encoded enzyme located in the mitochondrial matrix. As the first step in the Tricarboxylic Acid (TCA) cycle, the activity of this enzyme is often used as an indicator of cellular oxidative capacity. Citrate synthase activity per unit of mitochondrial protein increased across development in all genotypes (Fig. 3) (mito-nuclear genotypes: instar, *P*< 0.0001; natural genotypes: instar, *P* < 0.0001). There were also genotype-specific effects on citrate synthase activity. The energetically compromised (*simw^501^*);*OreR* genotype had elevated citrate synthase activity relative to other genotypes across all three instars (Fig. 3), resulting in significant epistatic mito-nuclear variance for this measure of oxidative capacity (mito x nuclear, *P* = 0.022) (Supplemental Table 3). Genotype-by-instar interactions significantly affected citrate synthase activity in the natural genotypes (instar x genotype, *P* = 0.010). Genotypes could be categorized as those for which citrate synthase reaches is maximal level by the second instar (e.g., *VT10* and (*ore*);*Aut*) and those for which second-instar mitochondria have citrate synthase activity levels intermediate to first- and third-instar mitochondria (e.g., *VT4* and *(ore*);*OreR*) (Fig. 3).

**Figure 3.**
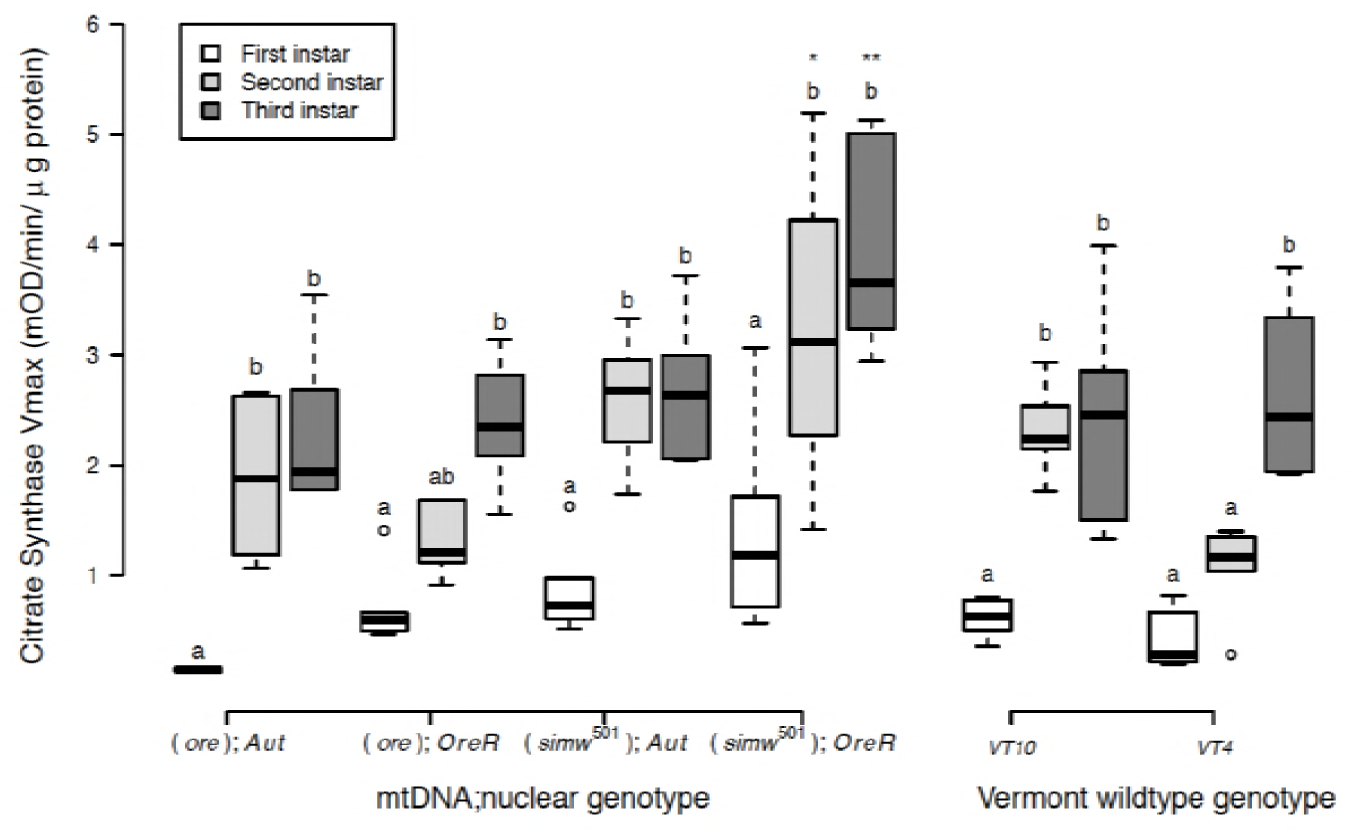
Citrate synthase activity. Oxidative capacity, measured by citrate synthase activity (V_max_) per unit of mitochondrial protein, increased significantly across instars and was largest in the energetically-compromised (*simw^501^*);*OreR* genotype. A) While all mito-nuclear genotypes increased oxidative capacity throughout development, there was significant variation among genotypes. (*simw^501^*);*OreR* larvae had significantly higher oxidative capacity than its nuclear genetic control (*ore*);*OreR* in the second (\**P_Tukey's_* = 0.015) and third instars (\*\**P_Tukey's_* = 0.008). The *simw^501^* mtDNA had no effect in the *Aut* background (*P _Tukey's_* > 0.833 in both instars), resulting in a significant mtDNA x nuclear interaction (*P* = 0.022). Wild-type genotypes from Vermont also varied significantly in the extent to which oxidative capacity reached its maximal level in the second versus third instar of development (Supplemental Table 3). Different letters within genotypes denote significantly different means at *P _Tukey's_* < 0.006, and asterisks designate significant differences between genotypes of the same larval instar.

In addition to aerobic, oxidative ATP production, *D. melanogaster* larvae use anaerobic, glycolytic ATP production that results in the production of lactate. There was significant genetic variation in the extent to which larvae accumulated lactate during development. Second-instar larvae of some genotypes significantly accumulated lactate, while others genotypes did not accumulate any lactate across development (Fig. 4A). This pattern was observed in both the mito-nuclear genotypes (instar x mtDNA x nuclear, *P* = 0.033) as well as in the natural genotypes (instar x genotype, *P* = 0.009). The energetically compromised (*simw^501^*);*OreR* genotype accumulated the highest amounts of lactate in the second instar, relative to other genotypes, resulting in a strong mito-nuclear interaction (Fig. 4B and Supplemental Table 4). However, the natural genotype *VT4* also accumulated high levels of lactate in second-instar larvae (Fig. 4A). Furthermore, genotypes that had intermediate levels of citrate synthase activity during the second instar (e.g., *VT4* and *(ore*);*OreR*) also tended to have increased lactate accumulation during the second instar.

**Figure 4.**
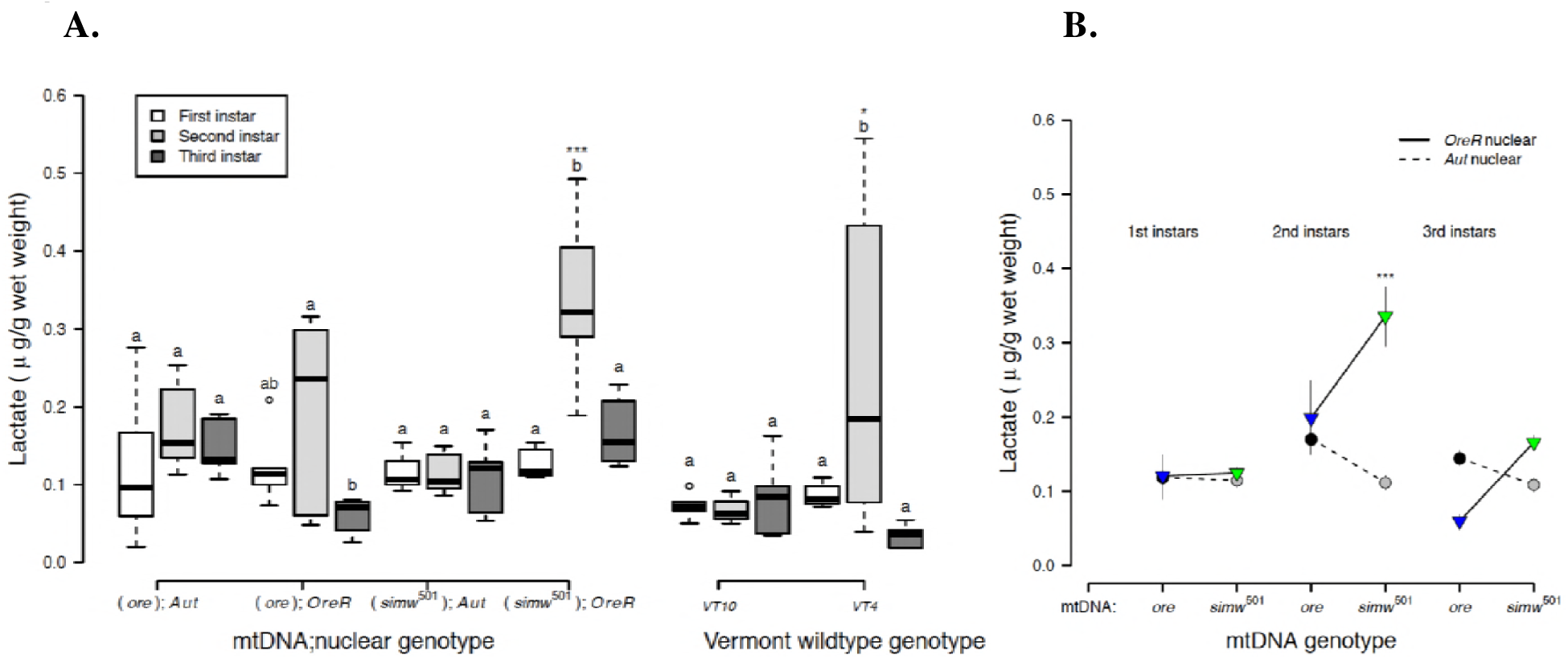
Lactate accumulation. A) Lactate levels per gram of larvae, varied significantly among genotypes in second-instar larvae and was highest in the energetically-compromised (*simw^501^*);*OreR* genotype. Genetic variation for second-instar larval lactate levels was also observed among wild-type genotypes from Vermont (instar x genotype, *P* = 0.009) (Supplemental Table 4), with *VT4* having significantly more lactate than *VT10* (\*\**P_Tukey's_* = 0.014). B) There was a significant instar x mtDNA x nuclear interaction effect on lactate levels (*P* = 0.033) (Supplemental Table 4). (*simw^501^*);*OreR* larvae had significantly higher lactate levels than all other genotypes in the second instar (\**P_Tukey's_* < 0.015). B) Different letters within genotypes denote significantly different means at *P _Tukey's_* < 0.036, and asterisks designate significant differences between genotypes of the same larval instar.

### The mito-nuclear incompatible genotype accumulates more ROS and has lower mitochondrial membrane potential

All genotypes had significantly increased levels of H_2_O_2_ by the third instar, relative to earlier instars (*P* < 0.0001) (Fig. 5A and Supplemental Table 5). However, the energetically-compromised (*simw^501^*);*OreR* genotype had significantly elevated levels of H_2_O_2_ in the second instar, both relative to other genotypes and to first- and third-instars of the same genotype. This resulted in a significant effect of the instar x mtDNA x nuclear interaction on levels of H_2_O_2_ (*P* < 0.0001) (Fig. 5B and Supplemental Table 5).

**Figure 5.**
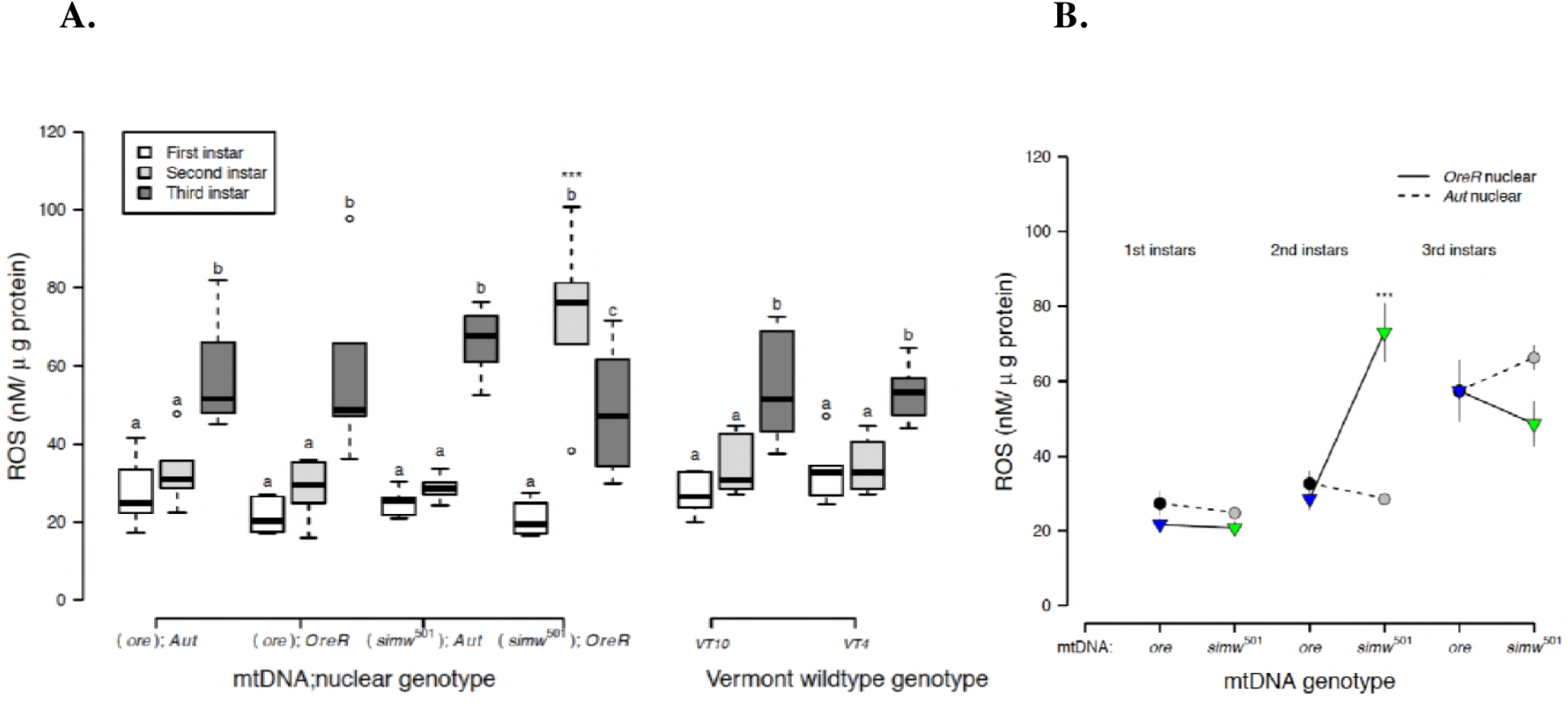
Reactive oxygen species (H_2_O_2_) A) ROS levels, measured as the concentration of H_2_O_2_ per gram of larvae, increased significantly across instars, and were highest in second-instar larvae of the energetically-compromised (*simw^501^*);*OreR* genotype. B) There was a strong effect of instar on ROS levels (instar, *P* = 2.347e-12), but this pattern varied among mito-nuclear genotypes (instar x mtDNA x nuclear, *P* = 5.166e-05) (Supplemental Table 5). Second-instar (*simw^501^*);*OreR* larvae had significantly higher ROS levels relative to all other genotypes (\*\*\**P_Tukey's_* < 0.0001), while all other genotypes had similar patterns of increasing ROS throughout development. The interaction between instar and genotype did not affect ROS levels among wild-type genotypes (Supplemental Table 5), which had a similar pattern to the control mito-nuclear genotypes. Different letters within genotypes denote significantly different means at *P _Tukey's_* < 0.041, and asterisks designate significant differences between genotypes of the same larval instar.

Because mitochondrial membrane potential (ΔΨ_m_) provides the driving force that is utilized by ATP synthase (complex V of OXPHOS) to make ATP and is used as an indicator of mitochondrial viability and cellular health, we tested whether this was disrupted in (*simw^501^*);*OreR*. All genotypes, except (*simw^501^*);*OreR*, maintained high levels of mitochondrial membrane potential in second- and third-instar larvae (Fig. 6A and Supplemental Table 6). The energetically-compromised (*simw^501^*);*OreR* genotype had significantly lower mitochondrial membrane potential relative to other genotypes in both second- and third-instar larvae. The effect of the mito-nuclear interaction on this marker of mitochondrial and cellular health was particularly pronounced in the second instar (instar x mtDNA x nuclear *P* < 0.0001) (Fig. 6B and Supplemental Table 6).

**Figure 6.**
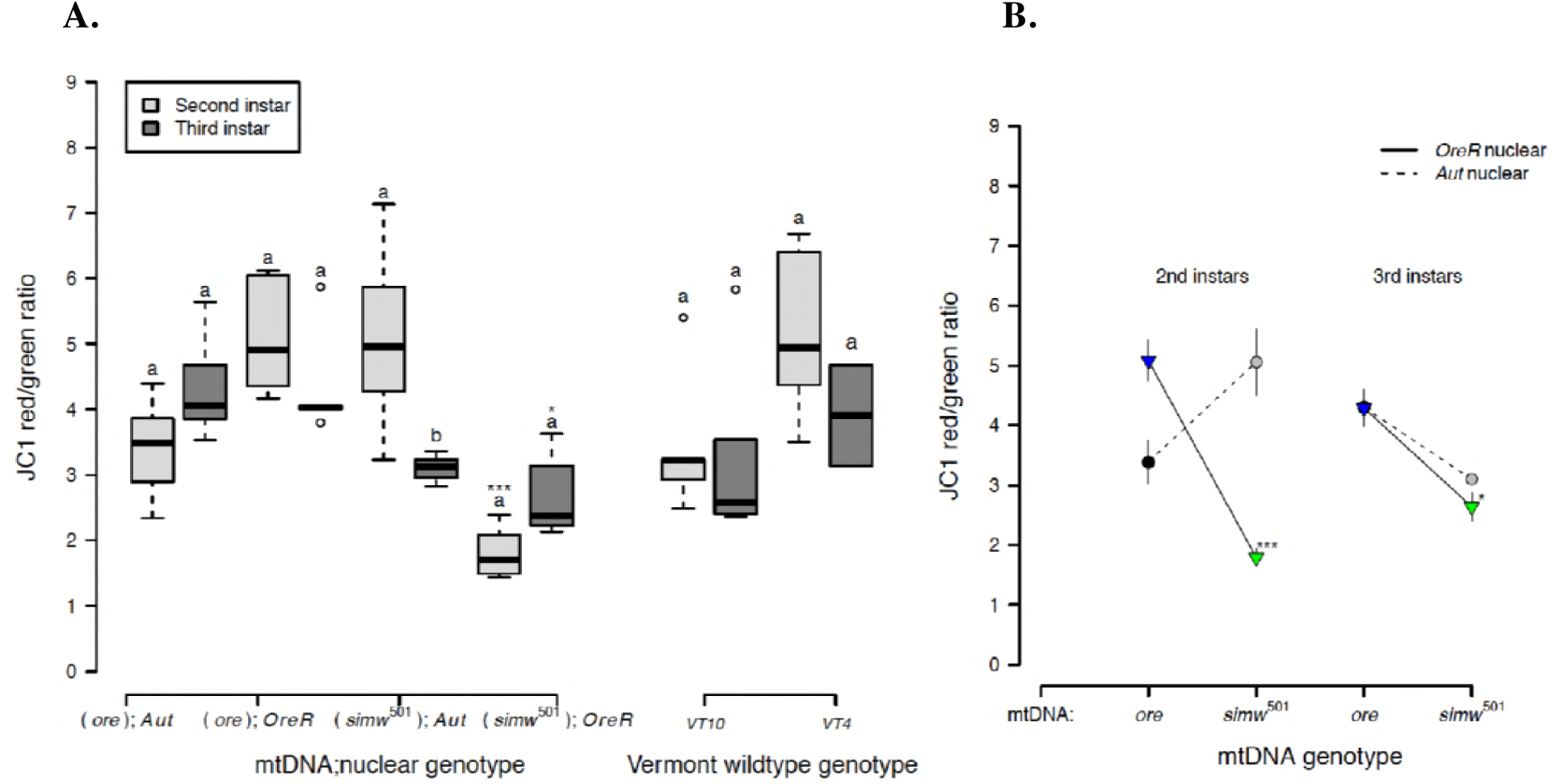
Mitochondrial membrane potential (ΔΨ_m_) The energetically compromised (*simw^501^*);*OreR* genotype had significantly decreased mitochondrial quality, as measured by the mitochondrial membrane potential (ΔΨ_m_). A) Both mito-nuclear and wild-type genotypes from Vermont generally maintained high membrane potential in second- and third-instar larvae. However, (*simw^501^*);*OreR* larvae had significantly lower mitochondrial membrane potential than the nuclear genetic control (*ore*);*OreR* in the second (\*\*\**P_Tukey's_* < 0.0001) and third instars (\**P_Tukey's_* = 0.016). B) This effect of the *simw^501^* mtDNA was not evident in the *Aut* background, where it increased membrane potential in second-instar larvae and had no effect in third-instar larvae (*P_Tukey's_* = 0.167). This resulted in a significant effect of an instar x mito x nuclear interaction (*P* = 1.580e-05) (Supplemental Table 6). Values above 2 typically indicate healthy mitochondria. Different letters within genotypes denote significantly different means at *P _Tukey's_* < 0.002, and asterisks designate significant differences between genotypes of the same larval instar.

In summary, the maintenance of mitochondrial respiration in the energetically compromised (*simw^501^*);*OreR* genotype across second and third instars was coincident with significant increases in oxidative capacity of mitochondria, increased lactate and ROS production during the second instar, and decreased mitochondrial membrane potential.

## Discussion

### Ontogenetic shifts in the relationship between metabolic rate and mass

Metabolic rates scale allometrically with mass, but the parameters that define this relationship vary among taxa, genotypes, life stages and environments (Glazier, 2005; Greenlee *et al.*, 2014). We found that the relationship between mass and metabolic rate differed significantly among larval instars of *D. melanogaster*. Metabolic scaling in developing animals has been described as an “impasse of principles,” wherein the basic tenant of metabolic allometry—that the physiological principles of organisms are relatively conserved—is at odds with the basic tenant of development—that the physiological state of organisms is dynamic across ontogeny (Burggren, 2005). Insect development involves complex changes in cellular energy demand and body composition that likely affect how metabolic rate scales with mass. Thus, models and principles of interspecific allometric scaling may not be applicable to ontogenetic scaling.

We observed a shift from hypermetric scaling in first-instar larvae (*b*>1), to isometric allometry in second-instar larvae (*b*=1), followed by hypometric allometry in third-instar larvae (*b*<1). The shift in metabolic scaling towards lower mass-specific metabolic rates in larger instars, was in spite of our observation that larger instars had seemingly greater capacity for oxygen-dependent ATP production, as indicated by increased levels of citrate synthase activity per unit of mitochondria. Nevertheless, mitochondrial oxygen consumption linked to ATP production was maintained at similar levels across second- and third-instar larvae. These patterns suggest that although there may be increased oxidative capacity of mitochondria as development progresses, mitochondrial respiration and organismal respiration are not simple reflections of oxidative capacity, but rather are emergent properties of organellar, cellular and organismal processes.

The ontogenetic change in metabolic scaling that we observed may reflect a change in energy demand across development as larval growth transitions from cell proliferation to cell growth. Hypermetric metabolic scaling exponents (b>1), where metabolic rates of larger individuals are greater per unit mass, could result from the increased energetic costs associated with the rapid cell proliferation and increase in cell number early in *Drosophila* development (O’Farrell, 2004; Vollmer *et al.*, 2017). Later in development, larval accumulation of mass occurs primarily via increases in cell volume (O’Farrell, 2004), reducing the surface area to volume ratio of cells and potentially limiting metabolism. These observations support studies, collectively grouped under Resource Demand (RD) models, that suggest that metabolic scaling is driven by an intrinsic metabolic demand from the cellular level to tissue growth potential (VonBertalanffy and Pirozynski, 1953; Shin and Yasuo, 1984, 1993; Ricklefs, 2003; Glazier, 2005). In this way, an organism's metabolic rate (and by extension metabolic scaling) across development is a reflection of the potential of tissues for proliferation and growth (Ricklefs, 2003). Our observations also contribute to a small, but growing number of insect studies that support a conceptual framework for patterns of intraspecific ontogenetic scaling where completion of growth in holometabolous insects is correlated with decreased mass-specific metabolic rates (Glazier, 2005). In both the tobacco hornworm *Manduca sexta,* and the silkworm *Bombyx mori*, metabolic scaling exponents also decrease across ontogeny (Blossman-Myer and Burggren, 2010; Callier and Nijhout, 2011, 2012; Sears *et al.*, 2012).

Metabolic scaling with mass may also be influenced by the biochemical composition of the body. System Composition (SC) models hypothesize that ontogenetic changes in metabolic scaling reflect shifts in body composition and the relative proportions of metabolically active versus inert or “sluggish” tissues (Glazier, 2005; Isler and VanSchaik, 2006; Greenlee *et al.*, 2014). Tissue lipid composition and lipid storage change across development in *Drosophila*, with a net increase of metabolically-inert storage lipids like triacylglycerides across development (Carvalho *et al.*, 2012). The increased contribution of less metabolically active tissue to the body could also contribute to the observed developmental shift from hypermetric to hypometric metabolic scaling. In *Manduca*, the contribution of metabolically active gut tissue to the body decreases across development, which may contribute to an increasingly hypometric metabolic scaling with mass across development (Callier and Nijhout, 2012).

Genetic variation in body composition across development could also underlie the significant genetic variation in metabolic scaling that we observed in third-instar larvae. If genotypes differ in the degree to which they accumulate mass in the third instar via different types of energy storage, this could generate genetic variation for how metabolic rate scales with mass. Midway through the third instar, *D. melanogaster* membrane-lipid accumulation is paused, while levels of storage lipids like triacylglycerides increase (Carvalho *et al.*, 2012).This suggests a transition in the third instar from metabolism supporting membrane synthesis and cell proliferation to metabolism supporting mass accumulation via lipid storage. If genotypes vary in the timing or extent of this switch, as we have seen for mitochondrial metabolism (described below), this could contribute to the greater genetic variation for metabolic rate and scaling that we observed in this developmental stage.

### Genetic variation in cellular metabolism may support similar organismal outcomes

Aerobic organisms can generate ATP via mitochondrial OXPHOS, but also anaerobically via glycolytic pathways that are supported by fermentation-generated NAD^+^ (e.g., by lactate production). We observed a consistent pattern across genotypes of increased oxidative capacity to aerobically produce ATP across development, consistent with a metabolic switch from glycolytic to mitochondrial production of ATP regulated by the *Drosophila* estrogen-related receptor *dERR* (Tennessen *et al.*, 2011; Tennessen and Thummel, 2011). However, genotypes differentiated into two categories of metabolic phenotypes—those that appeared to switch to mitochondrial ATP production before the second instar and accumulated very little lactate, and those that appeared to rely on glycolytic ATP production with significant lactate accumulation during the second instar. Yet, despite this genetic variation in how second-instar larvae were generating ATP, the organismal metabolic rate of second-instar larvae was more robust to genetic variation than were the metabolic rates of other instars. We also observed that despite this developmental switch from glycolytic to mitochondrial ATP production, *in vitro* mitochondrial respiration rates per unit mitochondrial protein remained constant across second- and third-instar larvae. Again, this highlights that organellar and organismal metabolic rates emerge from cellular, tissue and organismal-level processes, and are not simple reflections of the underlying metabolic pathways being used. In this way, higher levels of biological organization may buffer and potentially shelter genetic variation in metabolism from selection.

Using glycolytic ATP production may seem counterintuitive and less efficient than oxidative ATP production. However, glycolytic ATP production may provide several developmental advantages. First, it meets the bioenergetic needs of growth by providing abundant ATP. Despite the low yield of ATP per glucose consumed, the percentage of total cellular ATP produced from glycolysis can exceed that produced by OXPHOS (Lunt and Vander-Heiden, 2011). Second, there are reports of moonlighting functions of glycolytic enzymes translocating to the nucleus where they act as transcription factors to promote proliferation (Marden, 2013; Lincet and Icard, 2014; Boukouris *et al.*, 2016). Third, glycolytic ATP production enables flux through the pentose phosphate and TCA pathways to provide carbon-backbone intermediates for building macromolecules such as ribose sugars for DNA, amino acids for proteins, glycerol and citrate for lipids, as well as reducing power to support cell proliferation and growth during development (Tennessen *et al.*, 2011, 2014; Tennessen and Thummel, 2011; Lunt and Vander-Heiden, 2011).

*dERR* is responsible for a vital transcriptional switch of carbohydrate metabolism in second-instar larvae (Tennessen *et al.*, 2011) that coincides with increases in lactate dehydrogenase (*dLDH*) and lactate accumulation (Li *et al.*, 2017). dLDH activity recycles NAD^+^ which allows for continued glycolytic ATP production and supports the TCA cycle in generating cellular building blocks. Furthermore, dLDH expression and lactate production results in the accumulation of the metabolic signaling molecule L-2-hydroxyglutarate that affects genome-wide DNA methylation and promotes cellular proliferation (Li *et al.*, 2017). We found that second-instar lactate accumulation was strongly affected by genotype, suggesting differential timing of this switch among both wild-type and mito-nuclear genotypes. Investigating potential bioenergetics and life-history consequences of this genetic variation may reveal whether different metabolic strategies at the sub-cellular level fund similar or distinct fitness outcomes at the organismal level. This is critical for understanding whether populations harbor genetic variation in biochemical pathways that ultimately has similar fitness outcomes or whether we should expect to see the signatures of selection acting on enzymes that control shifts in metabolic flux (e.g., Flowers *et al.*, 2007; Pekny *et al.*, 2018).

### Physiological compensation in a mito-nuclear incompatible genotype comes at a cost

Mitochondrial respiration coupled to ATP production was maintained in the mito-nuclear incompatible genotype at *in vitro* levels similar to control genotypes, despite compromised OXPHOS via a presumed defect in mitochondrial protein synthesis in this genotype (Meiklejohn *et al.*, 2013). The maintenance of mitochondrial respiration in this genotype was accompanied by increases in mitochondrial oxidative capacity, measured by citrate synthase activity, and glycolytic ATP production, measured by lactate accumulation, relative to control genotypes. These increases may reflect physiological compensation to maintain ATP levels in a genotype whose mitochondria consume similar levels of oxygen but are less efficiently generating ATP. We suggest that by using the functional complementation of both glycolytic and mitochondrial ATP production, this genotype is able to synthesize the ATP needed to support its development.

Physiological compensation can have diverse and sometimes counter-intuitive costs paid over the lifespan that can adversely affect fitness. While (*sim^w501^*); *OreR* appears able to physiologically compensate to survive larval development, this genotype has delayed development and compromised pupation height, immune function and female fecundity (Meiklejohn *et al.*, 2013; Zhang *et al.*, 2017; Buchanan *et al.*, 2018). Additionally, while *in vitro* mitochondrial respiration in this genotype was maintained similar to other genotypes, larval metabolic rate in this genotype was elevated, potentially via compensatory upregulation of aerobic capacity to supply ATP (Hoekstra *et al.*, 2013). Thus, even when drawing on both glycolytic and oxidative ATP production, individuals with this mito-nuclear incompatibility may produce energy supplies very close to demand during larval growth. Previous results from our lab support this model; when development of (*sim^w501^*); *OreR* was empirically accelerated, development and reproduction were even more compromised, suggesting that this genotype has limited capacity to compensate the defect in OXPHOS (Hoekstra *et al.*, 2013, 2018). This genotype may use all of its aerobic scope to complete normal development compared to the other genotypes, leaving few resources leftover for other aspects of fitness. Once the demands of growth are removed, this genotype appears to regain some aerobic scope, as larvae that survived to pupation also completed metamorphosis and had normal adult size and metabolic rates (Hoekstra *et al.*, 2013, 2018). However, the costs paid out during development appear to have significant impacts on adult fecundity. Both female and male fecundity were severely compromised in this genotype when development occurred at warmer temperatures that increase biological rates and energy demand (Hoekstra *et al.*, 2013; Zhang *et al.*, 2017).

At the cellular level, physiological compensation in (*sim^w501^*); *OreR* larvae may be a source of oxidative stress, indicated by higher levels of H_2_O_2_, relative to other genotypes. H_2_O_2_ is a byproduct of the mitochondrial electron transport system (ETS) that supports OXPHOS in healthy cells, and we observed increases in H_2_O_2_ as oxidative capacity increased across development in all genotypes. However, compromised electron flow through the ETS can increase H_2_O_2_ levels and generate oxidative stress (Somero *et al.*, 2017). There are two ways that this may be occurring in (*sim^w501^*); *OreR* mitochondria. First, upregulation of the TCA cycle to supply more NADH for ATP production via the ETS may increase production of superoxide anion at Complex I. Second, there may be stoichiometric imbalance in the ETS due to presumably normal levels of cytoplasmically-translated Complex II but compromised levels of the mitochondrially-translated downstream OXPHOS complexes in this genotype. This could result in backflow of electrons that can produce superoxide ions when the ratio of reduced:unreduced coenzyme Q become elevated. The idea that (*sim^w501^*); *OreR* individuals are experiencing oxidative stress suggests an alternative interpretation of the elevated citrate synthase activity that we observed in this genotype. Levels of citrate synthase were increased in the blue mussel *Mytilus trossulus* in response to heat stress, a change that was coupled with increases in isocitrate dehydrogenase (IDH), which generates NADPH^+^ to support H_2_O_2_-scavenging reactions in the mitochondria (Tomanek and Zuzow, 2010). This highlights the importance of considering that TCA cycle enzymes provide important functions beyond their role in OXPHOS, as they provide substrates for biosynthesis, support antioxidant reactions, and act as signaling molecules (Marden, 2013; Boukouris *et al.*, 2016; Somero *et al.*, 2017).

Finally, we observed that (*sim^w501^*); *OreR* mitochondria could support mitochondrial oxygen consumption linked to ATP production at wild-type levels despite the fact that their membrane potential was significantly reduced. There is precedence for this observation. For example, mitochondrial diseases with OXPHOS defects are correlated with a suite of metabolic phenotypes that include upregulated glycolysis, lactate accumulation, elevated ROS, and decreased mitochondrial membrane potential, but stable ATP levels (Szczepanowska *et al.*, 2012; Frazier *et al.*, 2017). ROS act as essential secondary messengers in cellular homeostasis, but above a certain threshold level can be dangerous and lead to apoptosis (Giorgio *et al.*, 2007; Bigarella *et al.*, 2014). A potential regulatory and defense mechanism is to decrease the mitochondrial membrane potential (e.g., by uncoupling) to reduce further ROS production and protect the cell from oxidative damage (Dlasková *et al.*, 2006). Our data cannot distinguish in this mito-nuclear genotype whether upregulation of citrate synthase and decreased membrane potential in the mitochondria are the cause or the consequence of oxidative stress. However, new models from ecophysiology (Tomanek and Zuzow 2010), developmental physiological genetic (Tennessen *et al.* 2011, 2014; Li *et al.* 2017) and disease (Ward and Thompson 2012) systems provide promising paths for future elucidation of the mechanisms by which mitochondrial-nuclear genetic variation scales up to organismal fitness variation.

In conclusion, the dramatic and rapid growth of *Drosophila* during ontogeny requires a precise and genetically determined metabolic program that enhances biosynthesis and proliferation coupled with a tight temporal coordination. Here, we have shown how genetic variation influence patterns of metabolism in both natural and mito-nuclear genotypes of *Drosophila* during its developmental progression. Our study reveals that genetic defects in core physiology can be buffered at the organismal level via physiological compensation and that natural populations likely harbor genetic variation for distinct metabolic strategies in development that generate similar organismal outcomes.

## Acknowledgements and funding

We would like to thank Madeleine Koenig who was supported by the Undergraduate Creative Activities and Research Experience (UCARE) program at UNL for her assistance with sample preparation. The research was supported by NSF-IOS CAREER Award 1149178, NSF EPSCoR Track II Award 1736249, and funds from the University of Nebraska-Lincoln to OBM, CRJ and KLM.

